# Systematic interrogation of the distinct metabolic potential in gut microbiomes of inflammatory bowel disease patients with dysbiosis

**DOI:** 10.1101/640649

**Authors:** Almut Heinken, Ines Thiele

## Abstract

The human gut microbiome plays an important role in human health. In order to investigate changes in metabolic activity associated with dysbiotic microbiomes, we retrieved strain-level relative abundances from metagenomics data from a cohort of pediatric Crohn’s Disease patients with and without dysbiosis, and healthy control children to construct a personalized microbiome model for each sample using the AGORA resource of genome-scale gut microbial reconstructions. Subsequently, we systematically profiled each individual microbiome by predicting the quantitative biosynthesis potential for all secreted metabolites as well as the strain-level contributions to each metabolite in each individual microbiome. The predicted fecal metabolomes and strain-metabolite contributions of microbiomes from patients with dysbiosis were distinct from healthy controls and patients without dysbiosis. Finally, we validated the predicted amino acid production profiles against fecal metabolomic measurements. Taken together, we presented an efficient, scalable, tractable computational approach to systematically interrogate the metabolic potential of individual microbiomes.

## Introduction

The human gut microbiome plays an important role in human health and disease. It performs important functions, such as maturation of the host immune system, digestion of food, synthesis of short-chain fatty acids, vitamins, and amino acids, and protection against pathogens (Shafquat et al., 2014; Tremaroli and Bäckhed, 2012). Changes in microbiome composition have been linked to complex multifactorial diseases, e.g., type 2 diabetes, metabolic syndrome, and nonalcoholic fatty liver syndrome (Okubo et al., 2018; Tremaroli and Bäckhed, 2012). Another multifactorial disease with unclear etiopathology is inflammatory bowel disease (IBD), which has also been linked to an altered gut microbiome composition (Sheehan et al., 2015). IBD consists of two subtypes, ulcerative colitis (UC) and Crohn’s Disease (CD), for which different changes in gut microbial composition have been reported (Sheehan et al., 2015). It remains unclear whether gut microbial dysbiosis is a cause or consequence of the disease (Knox et al., 2019) highlighting the need for a mechanistic understanding of the role of the microbiome in IBD. Suggested factors contributing to the pathogenesis of IBD include genetics, diet, lifestyle, and the gut microbiome (de Souza et al., 2017). The complexity of the factors contributing to IBD means that personalized treatment approaches are likely to be required (Knox et al., 2019). Hence, there is an urgent need to elucidate the links between these factors and the pathogenesis of IBD, and ultimately to propose potential therapeutics targeting the diet-host-microbiome axis (Knox et al., 2019).

A number of studies have reported differences in the abundances of certain taxa between IBD patients and healthy controls, identified through 16S rRNA sequencing or metagenomic approaches (Gevers et al., 2014; Knox et al., 2019; Lewis et al., 2015). A decrease in Bacteroidetes and Firmicutes taxa, especially anti-inflammatory genera, such as *Faecalibacterium* and *Roseburia*, and a corresponding increase in Proteobacteria (e.g., *Escherichia coli*) and Fusobacteria taxa has been consistently reported (Knox et al., 2019). However, metagenomic approaches alone are insufficient to infer the functional metabolic activity of the microbiome (Knox et al., 2019). Thus, functional, pathway-based analyzes are required to elucidate not only the changes in composition in the gut microbiomes of IBD patients but also the metabolic changes that could serve as a target for therapeutic interventions.

A useful approach to gain insight into the metabolic activity of a system is metabolomic measurements since metabolite profiles are a readout of what is happening at the biochemical level (Wishart, 2016). Several studies have analyzed the blood and/or fecal metabolome in IBD patients and cohorts (Franzosa et al., 2019; Marchesi et al., 2007; Santoru et al., 2017). For instance, IBD patients have reduced fecal levels of the short-chain fatty acid butyrate (Machiels et al., 2014; Marchesi et al., 2007). Fecal medium-chain fatty acids (e.g., pentanoate and hexanoate) were also reduced in IBD patients (De Preter et al., 2015). Increased fecal levels of amino acids and amines have been reported for IBD patients (Ni et al., 2017; Santoru et al., 2017), while fecal vitamin B levels have been reduced (Santoru et al., 2017). Amino acids, amines, and carnitines were increased in serum of pediatric IBD patients (Kolho et al., 2017). Perturbations of tryptophan metabolism have been reported for IBD patients (Nikolaus et al., 2017). There is an increasing evidence that these changes in patients’ metabolite pool can be partially linked to the microbiome. For instance, gut microbes produce short-chain fatty acids (den Besten et al., 2013), synthesize B and K vitamins (Shafquat et al., 2014), metabolize tryptophan into tryptamine, indole, and other degradation products (Agus et al., 2018), and transform primary into secondary bile acids (Ridlon et al., 2016). By performing metabolomics and metagenomic analyzes for the same fecal samples, it became possible to infer positive and negative correlations between species abundances and specific metabolites for cohorts of IBD patients and controls (Franzosa et al., 2019; Santoru et al., 2017). However, the mechanisms underlying these correlations remain unclear (Franzosa et al., 2019) and it is difficult to disentangle the contributions of microbial and host metabolism to altered metabolite levels.

To gain insight into the links between microbial and metabolic changes in the context of IBD, an integrative systems biology approach is necessary (de Souza et al., 2017). Such a computational systems biology approach can integrate omics data (e.g., metagenomics, metabolomics, and metatranscriptomics) as well as dietary information into a network “interactome” model that could propose potential disease mechanisms, biomarkers, and personalized therapies (de Souza et al., 2017). These computational modelling approaches may be constructed top-down by computationally inferring metabolite-microbe associations or bottom-up by linking metabolites and microbes manually based on biological knowledge. In the second case, computational models can provide testable mechanism-based hypotheses aiming at explaining metabolite-microbe associations. One such bottom-up systems biology approach is constraint-based reconstruction and analysis (COBRA) (Palsson, 2006). Briefly, COBRA relies on a manually curated genome-scale reconstruction of metabolism of a target organism, which can be converted into a mathematical model and subsequently interrogated through simulations, using established methods, such as flux balance analysis (O’Brien et al., 2015). COBRA models can be readily contextualized by implementing different types of data as constraints, e.g., metagenomics (Baldini et al., 2018), metabolomics (Aurich et al., 2016), proteomics (O’Brien and Palsson, 2015), or dietary information (Noronha et al., 2019). A suite of tools maintained by experts on constraint-based modelling, the COBRA Toolbox, is freely available for these purposes (Heirendt et al., 2019). Applying the COBRA approach to model host-microbe interactions uses ideally strain-resolved genome-scale reconstructions that represent the diversity of the human gut microbiome. To address this need, we have assembled a resource of 818 curated genome-scale reconstructions of human gut microbe strains, AGORA (Magnusdottir et al., 2017; Noronha et al., 2019). AGORA enables the creation of personalized microbiome models from metagenomics data (Baldini et al., 2018) and has valuable applications in studying microbe-microbe and host-microbe interactions (Magnusdottir and Thiele, 2018). AGORA has been successfully applied to investigating host-microbe interactions in colorectal cancer (Hale et al., 2018), predicting microbial cross-feeding at suboptimal growth (Henson and Phalak, 2018), modelling the lung microbiome in Cystic Fibrosis patients (Henson et al., 2019), predicting dietary supplements that promote short-chain fatty acid production in Crohn’s Disease patients (Bauer and Thiele, 2018), and analyzing microbial network patterns in relapsing Crohn’s Disease (Yilmaz et al., 2019).

Previously, Lewis et al. have performed metagenomic sequencing of the microbiomes of pediatric Crohn’s Disease patients and healthy control children and found that the Crohn’s Disease microbiomes stratified into two clusters, i.e., a “near cluster”, which resembled the healthy microbiomes in composition and a “far cluster” characterized by microbial dysbiosis that was distinct in composition from both healthy controls and the near cluster microbiomes (Lewis et al., 2015). We have previously constructed personalized models for a subset of 20 dysbiotic Crohn’s Disease and 25 healthy control microbiomes from this cohort (Heinken et al., 2019). We have then applied the COBRA approach to predict the bile acid deconjugation and biotransformation potential of each sample (Heinken et al., 2019). The computational modelling stratified the Crohn’s Disease microbiomes and healthy microbiomes by their bile acid metabolism profiles and revealed that the Crohn’s Disease microbiomes had a reduced potential to produce the secondary bile acid 12-dehydrocholate (Heinken et al., 2019). Moreover, the mechanistic modelling elucidated the exact strains contributing to the total bile acid deconjugation and biotransformation flux (Heinken et al., 2019). Thereby, we have established a computational framework to systematically explore the metabolic potential of a given gut microbiome with a metagenomic sample as the input.

Here, we substantially expanded the computation of metabolic profiles that previously only encompassed bile acids (Heinken et al., 2019) to a wide variety of metabolic subsystems. We systematically predicted the potential of each microbiome to secrete and take up all metabolites, for which biosynthesis pathways and transport reactions were present in the community models and identified the metabolites that best stratified the dysbiotic and non-dysbiotic individuals. For all secreted metabolites, we computed the respective contributing strains in each microbiome. Finally, we validated the predictions for amino acid metabolites against published metabolomic data from the same cohort. Taken together, we present a computational systems biology approach that bridges the gap between metagenomic and metabolomic data and provides mechanistic, testable hypotheses for metabolite-microbe associations.

## Results

We present a pipeline to predict the metabolic profile, i.e., the combined quantitative potential of all community members to take up dietary metabolites and secrete metabolic end products as well as the strain-level contributions to these overall fluxes, of a given individual microbiome (Figure 1, Methods). We comprehensively profile the metabolic potential of individual gut microbiomes of Crohn’s Disease microbiomes, with and without dysbiosis, and controls and stratified them according to these metabolic profiles. We demonstrate that the metabolite profiles of microbiomes from individuals with IBD and dysbiosis are distinct from both healthy microbiomes and IBD microbiomes with dysbiosis and identify the features that best separate these groups. Finally, we compare our results with published metabolomic data from the same individuals. In agreement with published findings, we predict that dysbiotic Crohn’s Disease microbiomes have an increased potential to synthesize amino acids.

**Figure 1:**
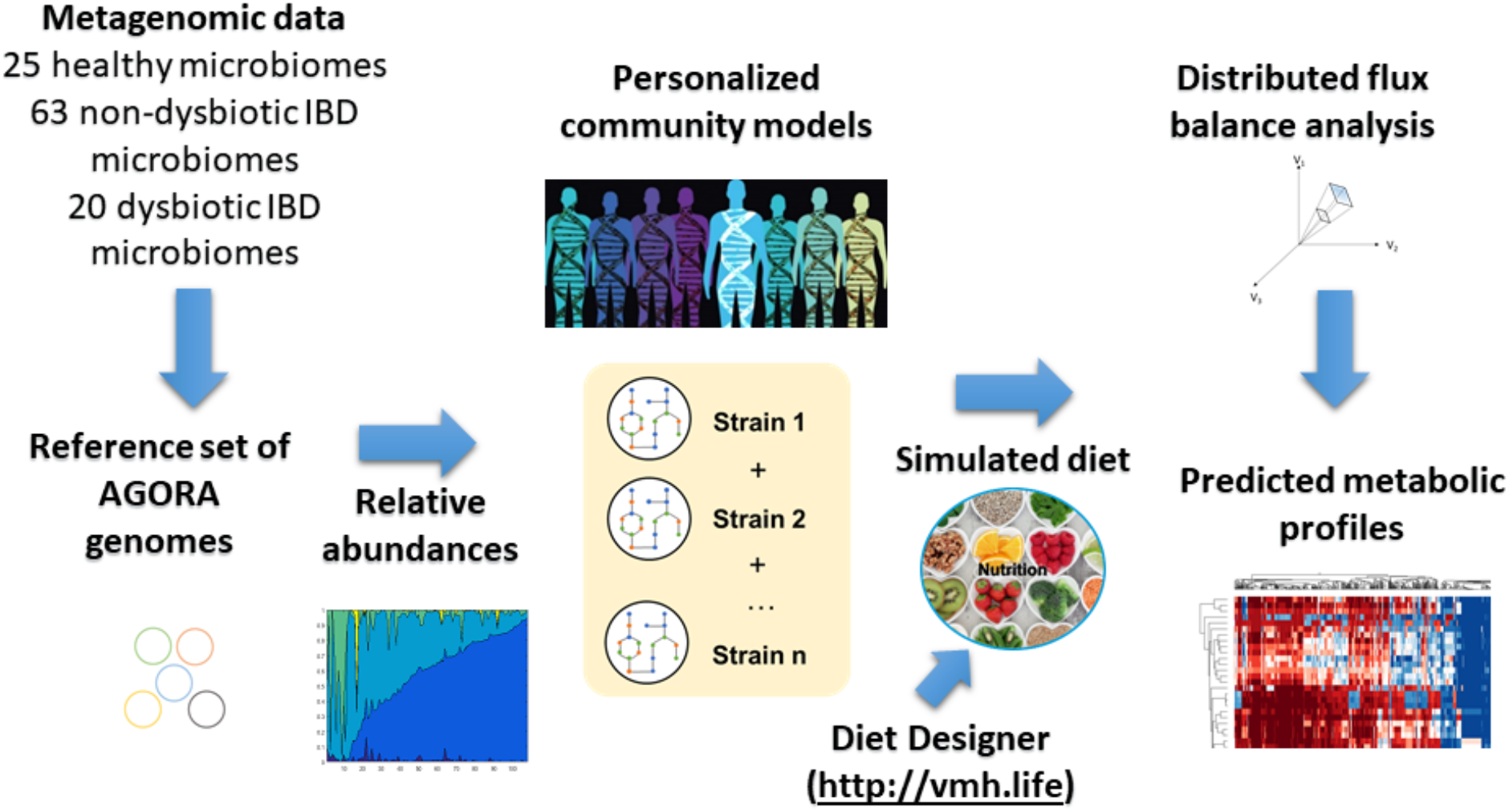
Schematic overview of the modelling framework established in this study. Metagenomic reads were mapped onto a reference set of AGORA (Magnusdottir et al., 2017) genomes, and the strain-level relative abundances were retrieved. For each metagenomic sample, a personalized microbiome model was constructed. The models were parameterized further with a simulated “Average European” diet retrieved from the Virtual Metabolic Human (Noronha et al., 2019) database. The metabolic profile of each microbiome model was then computed with distributed flux balance analysis (Heirendt et al., 2017).

### Profiling the range of metabolic capabilities present in the 108 microbiomes

We developed a large-scale, computationally efficient constraint-based modelling approach (Figure 1). First, we created a personalized microbiome model for each sample, which was constrained with a simulated Average European diet (Methods). The metabolic profile of each microbiome was then computed as follows: All dietary, fecal, and strain-specific exchange reactions present in the model were retrieved and the minimal and maximal fluxes through these exchange reactions were computed using distributed flux balance analysis (Heirendt et al., 2017) (Methods). This approach enabled a systematic evaluation of secretion potential, uptake potential, and strain-specific contributions for each metabolite that could be transported by at least one microbiome model.

Overall, the 108 microbiome models, responding 20 dysbiotic, 63 non-dysbiotic, and 25 control individuals, accounted for exchange reactions for 419 metabolites, for which the minimal and maximal fluxes were computed. Of these 419 metabolites, 143 could be secreted into the fecal compartment by at least one microbiome but were not taken up, i.e., they were of microbial origin but not dietary. Another 59 metabolites could only be taken up by at least one microbiome and were thus only present in the simulated diet. Finally, further 86 metabolites were both taken up and secreted (Table S1) meaning they were both dietary and microbial of origin. The remaining 131 metabolites that could be neither taken up nor secreted either lacked the necessary biosynthesis precursors in the given dietary input or were dead-end metabolites in the corresponding AGORA reconstructions. The 229 secreted metabolites belonged to diverse subsystems known to be present in the human gut microbiome, with amino acid metabolism (54 metabolites), and central energy metabolism (49 metabolites) being the biggest groups (Table S2).

Both the qualitative and quantitative potential to take up and secrete metabolites differed across the 108 microbiomes (Tables S1-S3). Some qualitative metabolite biosynthesis capacities were present in almost all microbiomes metabolite while others were rare (Table S1). In total, 116 secreted metabolites could be produced by 98 or more of the 108 microbiomes (>90%) but varied in their quantitative secretion potential (Figure S1, Table S1). Another 88 metabolites could be secreted by 27 to 96 microbiomes (25% to <90%). Finally, 25 metabolites were only produced by 17 or less microbiomes (<16%) and thus represented an uncommon biosynthesis capability. This latter set included methane (synthesized by 10 of 108 microbiomes, Table S1) highlighting that only few individuals carried methanogens in their microbiomes. Other less common fermentation products included valerate, butanol (12 microbiomes each), 1,2-propanediol (28 microbiomes), and 2,3-butanediol (31 microbiomes) (Table S1). Taken together, the systematic *in silico* metabolic profiling predicted that the analyzed microbiomes could convert the dietary inputs into a variety of metabolites from diverse subsystems known to be produced by human gut microbes and differed in their individual uptake and secretion profiles. The modelling framework captured the range of metabolic capabilities encoded by the human gut microbiome, and the variation in metabolic potential as a function of microbiome composition.

### Distinct metabolite uptake and secretion potential in dysbiotic compared with non-dysbiotic microbiomes

Since the microbial composition of the dysbiotic and non-dysbiotic individuals clearly differed (Lewis et al., 2015), we expected these microbial differences to be reflected in the predicted metabolic profiles. To test whether the computational modelling was able to stratify the dysbiotic and non-dysbiotic group, we performed a statistical analysis (Methods) of the predicted uptake and secretion fluxes. Of the 229 metabolites that were produced by at least one microbiome, 46 differed statistically significantly between the microbiomes of IBD patients and healthy controls (Wilcoxon rank sum test corrected for false discovery rate, Table S4a). Moreover, 121 metabolites had statistically significantly different production potential in the dysbiotic compared with the non-dysbiotic IBD cluster (Table S4b). The clearer separation between the dysbiotic and nondysbiotic IBD cluster than between healthy and IBD was in line with our expectations as the modelling framework was personalized with only the individuals’ gut microbial compositions and did not account for other factors (e.g., human metabolism and non-microbiome-mediated effects of medication). Taken together, dysbiotic IBD microbiomes were distinct in metabolite secretion potential as a direct consequence of their distinct microbial compositions.

Overall, the production potential for 106 metabolites differed between at least two of the three groups (Figure 2a-f). For instance, the dysbiotic IBD microbiomes were depleted in production potential for metabolites involved in glycan degradation, fermentation, and B-vitamin biosynthesis (Figure 2a-f). Metabolites with increased production potential in the dysbiotic cluster belonged mainly to the subsystems of amino acid metabolism, TCA cycle, simple sugars, and lipid metabolism (Figure 2a-f). We predicted an increased secretion potential in the dysbiotic cluster for 63 metabolites, including lactate, alanine, betaine, hydrogen sulfide, ethanol, ribose, and trimethylamine N-oxide (TMAO, Figure 2a-f). Increased fecal lactate has been reported for Crohn’s Disease patients (Franzosa et al., 2019). Another study, also in agreement with our predictions, reported increased fecal alanine in IBD patients (Santoru et al., 2017). Ribose has been found to be increased in serum of dogs with IBD (Minamoto et al., 2015). Hydrogen sulfide has been proposed to both worsen (Ijssennagger et al., 2016) and protect against (Wallace et al., 2018) gastrointestinal inflammation. On the other hand, a reduced secretion potential was predicted for 58 metabolites, among them being butyrate, nicotinamide, nicotinic acid, reduced riboflavin, and degradation products of mucins and other glycans (Figure 2a-f, Table S4b). Reduced SCFA levels, especially butyrate, are well-described for IBD (Marchesi et al., 2007). Decreased nicotinic acid levels have also been reported for IBD patients (Franzosa et al., 2019; Santoru et al., 2017). *Faecalibacterium prausnitzii*, which is well known to be depleted in Crohn’s Disease patients (Pascal et al., 2017), uses riboflavin as a redox mediator in an extracellular electron shuttle (Khan et al., 2012) explaining the decreased riboflavin reduction in the dysbiotic cluster. Riboflavin biosynthesis was also decreased in IBD microbiomes (Table S4a). Our results suggest a reduced functional capacity of the dysbiotic microbiomes to synthesize and secrete butyrate, glycan degradation products, and B-vitamins, and an increased capacity for production of certain energy metabolites (e.g., lactate), hydrogen sulfide, amino acids, and amines. To identify the secreted metabolites that best separated the three groups, a Random Forests analysis on the predicted net production fluxes was performed in MetaboAnalyst 4.0 (Chong et al., 2018). The metabolites found to best stratify the non-dysbiotic and dysbiotic IBD cluster were chorismate, L-lactate, phenol, D-ribose, phenylalanine, and nicotinamide (Figure 2g).

**Figure 2:**
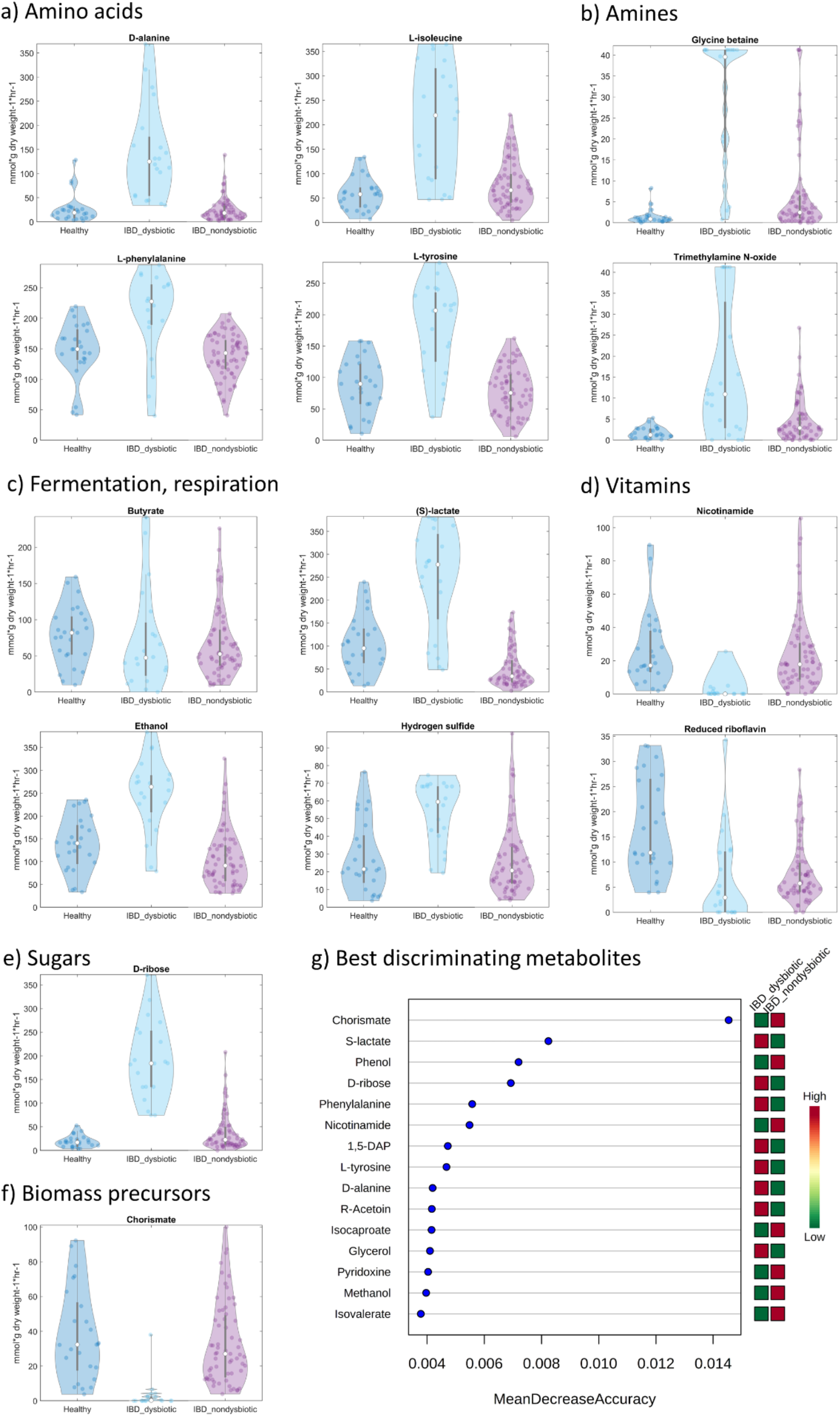
Predicted metabolite secretion profiles for the 108 microbiomes. a-f) Total metabolite secretion fluxes (mmol × person^−1^ × day^−1^) predicted for the microbiomes of 20 pediatric Crohn’s Disease patients with dysbiosis (IBD_dysbiotic), 63 pediatric Crohn’s Disease patients without dysbiosis (IBD_nondysbiotic), and 25 healthy control children (Healthy) for selected metabolites of interest. g) Random Forests analysis showing the secreted metabolites that best stratified the 20 dysbiotic and 63 dysbiotic IBD microbiomes.

As a next analysis, we computed the net metabolite uptake potential of each community and found it to be distinct in dysbiotic IBD microbiomes (Table S3, Table S4b). The computed net uptake potential represents the capacity of each individual gut microbiome to extract dietary nutrients and calories from the diet, which provides insight into the fate of dietary input. Overall, the uptake potential for 15 and 23 metabolites, respectively, differed between healthy and IBD and between the dysbiotic and nondysbiotic IBD cluster. The dysbiotic cluster was enriched in the potential to take up phenylalanine, leucine, pyridoxamine, nicotinic acid, nicotinamide, iron(ii), and several fatty acids but depleted in the potential to take up iron(iii), sulfate, lysine, and resistant starch (Table S4b). The reduced potential to metabolize resistant starch suggests a reduced capability to metabolize fiber into short-chain fatty acids, and indeed, butyrate production potential was decreased in the dysbiotic cluster (Figure 2a, Table S4b). On the other hand, an increased potential to take up B vitamins may indicate increased competition with the host over dietary B vitamins.

To link the predicted metabolite uptake and secretion potential to specific microbes, we calculated the Spearman correlation between metabolite secretion and uptake potential and species abundances. Strong correlations (>0.75) between species and metabolites were found for 66 secreted metabolites (Figure 3a) and for ten consumed metabolites (Figure S1). For instance, glycan degradation products strongly correlated with several *Bacteroides* spp., in agreement with *Bacteroides* being known glycan degraders (Porter and Martens, 2017) (Figure 3a). Secondary bile acids correlated with species known to biotransform bile acids (Figure 3a) as observed previously (Heinken et al., 2019). As expected, methane correlated positively with *Methanobrevibacter smithii* (Samuel et al., 2007), and p-cresol correlated with the known p-cresol producer *Clostridioides difficile* (Selmer and Andrei, 2001) (Figure 3a). *Ruminococcus bromii* and *Ruminococcus champanellensis* correlated with uptake of resistant starch and cellulose, respectively (Figure S1), in agreement with their known capability to degrade these fibers (Chassard et al., 2012; Ze et al., 2012). The remaining 163 secreted and 145 consumed metabolites did not strongly correlate with specific species. Thus, the uptake and production of these metabolites was carried out by a combination of multiple taxa.

**Figure 3:**
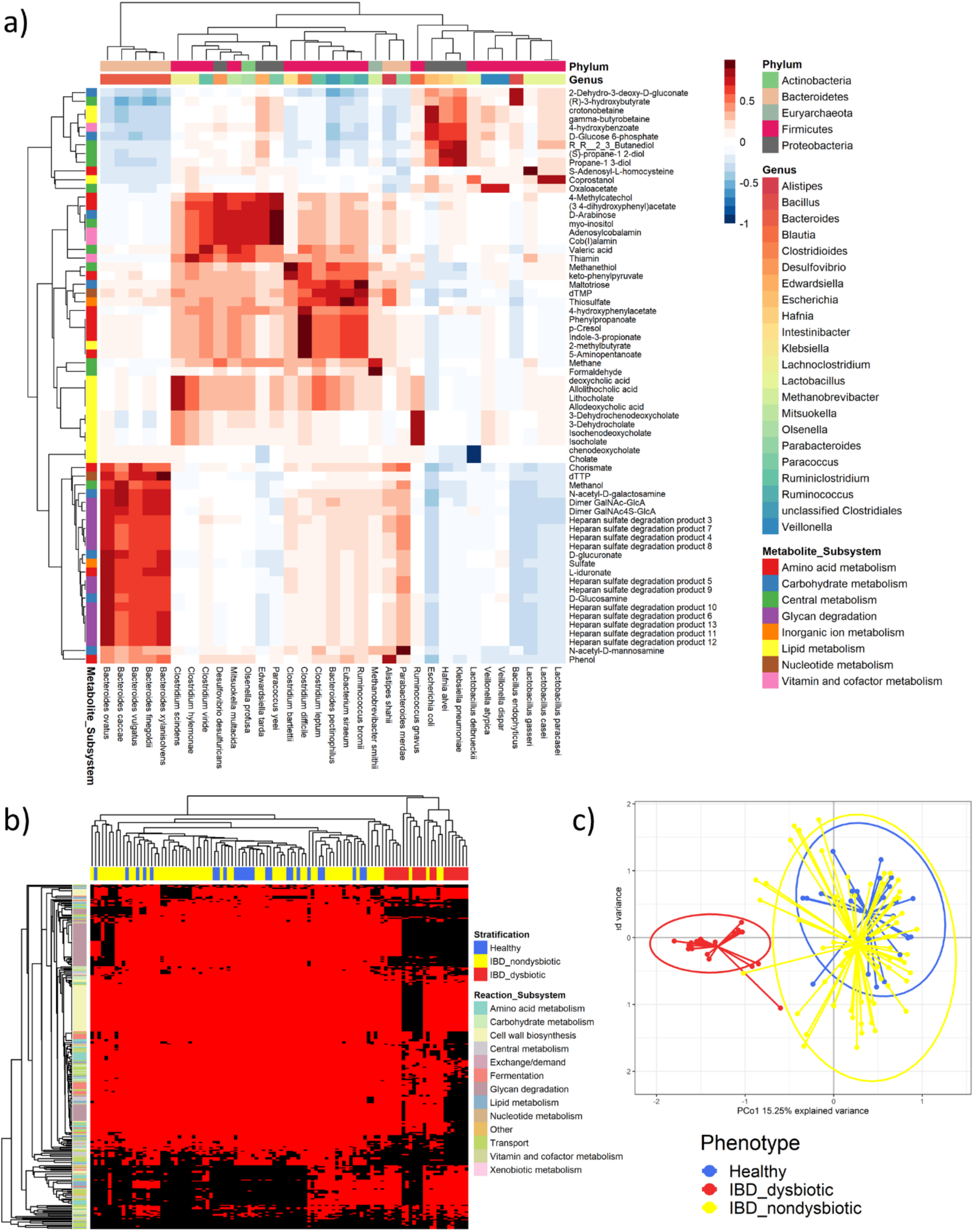
Metabolic properties and flux profiles computed for the 108 microbiomes. a) Spearman correlations between metabolite secretion fluxes (mmol × person^−1^ × day^−1^) and species-level relative abundances across all 108 microbiome models. Shown are only metabolites for which the positive or negative correlation with at least one species was higher than 0.75 or lower than −0.75, respectively. Rows show metabolites annotated by subsystem, and columns show species annotated by genus and phylum. b) Absolute reaction presence in the 108 microbiome models than differed significantly (p-value corrected for false discovery rate <0.05, Table S4b) between the 20 dysbiotic and 63 dysbiotic IBD microbiomes. Rows show reactions annotated by subsystem, and columns show microbiome models annotated by group. Red=reaction present, black=reaction absent. C) Principal components analysis of all 24,478 strain to metabolite contributions (mmol × person^−1^ × day^−1^, Table S5) computed for the 108 microbiome models.

### Absolute and quantitative presence of metabolic functions is altered in dysbiotic microbiomes

To explain the observed changes in metabolite uptake and secretion potential, we calculated and inspected the absolute presence of reactions in the microbiomes as well as the quantitative reaction abundances on the whole community, class, and genus level. The absolute presence of 52 and 312 reactions was distinct between healthy and IBD and between the dysbiotic and non-dysbiotic IBD cluster, respectively (Figure 3b, Table S4a-b). Thus, the dysbiotic IBD microbiomes were depleted or enriched in the absolute presence of certain pathways. Specifically, reactions involved in glycan degradation were absent in most dysbiotic microbiomes (Figure 3b).

The abundances of 454 reactions on the total community level, 3,621 reactions on the phylum level, and 30,173 reactions on the genus level differed significantly between healthy controls and IBD patients (Table S4a). Moreover, 1,374 reactions on the total community level, 5,944 reactions on the phylum level, and 42,580 reactions on the genus level were different in abundance between the dysbiotic and non-dysbiotic IBD cluster (Table S4b). Taken together, the microbiomes of dysbiotic IBD patients were distinct in the qualitative and quantitative presence of key reactions and pathways, which explains their aforementioned altered potential to consume and secrete metabolites. Thus, the microbial community structure and function was disturbed in dysbiotic states associated with IBD, as observed previously (Yilmaz et al., 2019), and suggests that changes in gut microbiome composition have a broad effect on the community’s metabolic potential.

### A wide variety of taxon to metabolite contributions are altered in IBD microbiomes

Fecal gut metabolite levels are altered in IBD patients, including many host-microbial co-metabolites, however, the contributions of specific microbes to these changes are often unclear (Franzosa et al., 2019). To gain insight into which microbial taxa are responsible for the altered metabolic profiles in dysbiotic IBD patients, we modelled the strain-to-metabolite contributions directly by predicting the quantitative contribution of each strain to each secreted metabolite in each individual microbiome.

All 601 strains present in at least one microbiome contributed to the secretion of at least one metabolite indicating they should be able to potentially influence host metabolism. In total, 24,478 strain-to-metabolite contributions were predicted, corresponding on average to 40.73 contributions per strain (Table S5). Of the 24,478 strain-to-metabolite contributions, 2,463 (10.06%) and 7,214 (29.47%), respectively, were statistically significantly different between healthy and IBD (Table S4a) and dysbiotic and non-dysbiotic IBD microbiomes (Table S4b). Hence, a wide variety of metabolic fluxes was distinct in dysbiotic microbiomes. A principal components analysis of all 24,478 strain-to-metabolite demonstrated that the dysbiotic IBD microbiomes clustered separately from both the healthy and the non-dysbiotic IBD microbiomes (Figure 3b). Again, this was expected as the models directly reflected the distinct microbiome compositions of the dysbiotic IBD patients.

Next, we investigated the contributions that were distinct in dysbiotic microbiomes by taxon. The contributions were summarized by phylum and by metabolite subsystem for the three clusters (Figure 4a). In the dysbiotic cluster, contribution flux profiles clearly differed from the healthy and non-dysbiotic IBD cluster, showing a drastic reduction in Bacteroidetes contributions accompanied by an increase in Proteobacteria and Fusobacteria contributions (Figure 4a). This result is in line with observations that the dysbiotic cluster was characterized by an increase in gammaproteobacterial genera, and a corresponding decrease in Bacteroidetes and Clostridia (Lewis et al., 2015). While the non-dysbiotic cluster’s phylum to subsystem contributions resembled overall the profile of the healthy cluster, there was a slight increase in Fusobacteria and Proteobacteria contributions compared with the latter (Figure 4a). This change may indicate that the individuals in the non-dysbiotic IBD cluster represent an early state in the development towards pronounced dysbiosis as observed in the dysbiotic cluster. The altered contribution profiles in the dysbiotic cluster were observed across all metabolic subsystems (Figure 4a) demonstrating a broad effect of microbial composition changes on metabolite fluxes.

**Figure 4:**
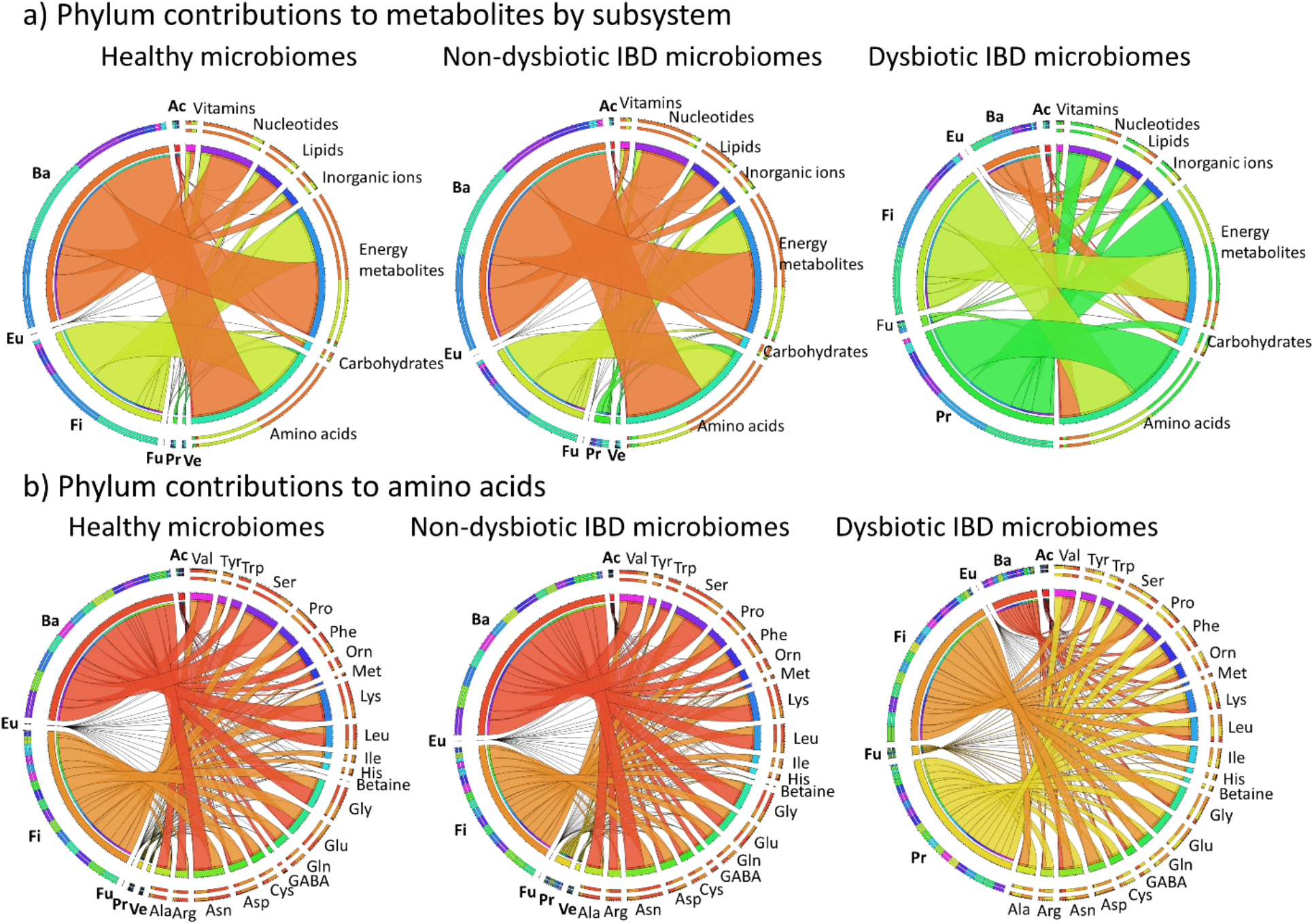
Strain to metabolite contributions computed for the three clusters summarized by the seven most prominent phyla. Shown are the contributions (mmol × person^−1^ × day^−1^, Table S5) summarized separately for the 25 healthy, 63 non-dysbiotic IBD, and 20 dysbiotic IBD microbiomes. a) Phylum-level contributions to all metabolites summarized by metabolite subsystem. b) Phylum-level contributions to 22 amino acid metabolites. Ac=Actinobacteria, Ba=Bacteroidetes, Eu=Eurarchaeota, Fi=Firmicutes, Fu=Fusobacteria, Pr=Proteobacteria, Ve=Verrucomicrobia. Phyla that did not contribute in a given cluster are omitted for that cluster.

To gain more insight into metabolites of interest that were enriched or depleted in the dysbiotic cluster, we extracted the strain contributions (Table S5) by metabolite, which is shown for the examples of butyrate, L-lactate, hydrogen sulfide, trimethylamine-N-oxide (TMAO), and nicotinamide in Figure S2-6. For instance, not only the total butyrate production flux was reduced in dysbiotic IBD microbiomes (Figure 2a) but the contributing strains also differed between dysbiotic and non-dysbiotic microbiomes (Figure S2). As expected, known butyrate producers, e.g., *Roseburia* spp., *Faecalibacterium praunitzii*, and *Eubacterium rectale*, contributed the majority of butyrate in healthy and non-dysbiotic IBD microbiomes (Figure S2). In dysbiotic microbiomes, butyrate secretion by these species was reduced (Figure S2). Reduced butyrate levels in IBD patients with dysbiosis have previously been attributed to reduced abundance of *Roseburia* and *Faecalibacterium* species (Machiels et al., 2014). For L-lactate, both the total production potential (Figure 2c) and the contributing strains (Figure S3) also varied between dysbiotic and non-dysbiotic microbiomes. In the microbiomes of healthy controls and in the non-dysbiotic cluster, L-lactate was mainly produced by Clostridia, e.g., *Eubacterium rectale*, *Anaerostipes hadrus*, and *Roseburia* spp. (Figure S3). In contrast, in the dysbiotic cluster, the main contributions came from Gammaproteobacteria genera (e.g., *Escherichia* and *Klebsiella*), lactic acid bacteria (*Lactobacillus*, *Enterococcus*, and *Streptococcus*), and *Veillonella* species (Figure S3). Similarly, hydrogen sulfide and TMAO were clearly enriched in the dysbiotic cluster (Figure 2b-c), which could be mainly attributed to gammaproteobacterial genera, such as *Escherichia*, *Klebsiella*, *Enterobacter*, and *Sutterella* (Figure S4-5). On the other hand, nicotinamide, which was clearly reduced in the dysbiotic cluster (Figure 2d), was synthesized by *Bacteroides* and *Parabacteroides* representatives (Figure S6). Thus, through metabolic modelling, we could retrieve a strain and individual-resolved, detailed snapshot of each metabolite of interest. Butyrate, L-lactate, TMAO, and hydrogen sulfide were produced by a combination of multiple species (Figures S2-5), explaining why no strong correlations with any individual species were observed for these metabolites in Figure 3a.

To summarize, we systematically interrogated the gut microbe-metabolite axis through constraint-based modelling. A reduction to metabolite contributions from commensal and beneficial taxa, such as *Roseburia* and *Faecalibacterium*, and a corresponding increase in contributions from potential detrimental taxa, such as *Escherichia*, was found for dysbiotic IBD microbiomes. For each metabolite, the exact contributing strains were identified (Figure S2-6, Table S5). Overall, commensal and beneficial taxa contributed more to metabolites that are thought to be relevant for health (e.g., butyrate, B-vitamins), and taxa associated with dysbiosis (i.e., Proteobacteria, Bacilli) contributed more to potentially harmful metabolites, such as lactate, hydrogen sulfide, and TMAO.

### Modelling provides evidence for increased proteobacterial amino acid biosynthesis potential in dysbiosis

It has been proposed that metabolic modeling could be used to link metagenomic and metabolomic findings (Heinken et al., 2019), which would require published metagenomic and metabolomic measurements form the same cohort. In a recent study by Ni et al. (Ni et al., 2017), fecal metabolomic profiles were indeed determined for the same cohort of healthy controls and pediatric Crohn’s Disease patients (Lewis et al., 2015) that we used in this study. The patients in the dysbiotic cluster had distinct fecal metabolomic profiles, which have been characterized by an increase in amino acids and amino acid derivatives (Ni et al., 2017). In agreements with these findings, we predicted an increased potential to synthesize 4-aminobutanoate (GABA), D-alanine, aspartate, betaine, glutamate, glycine, isoleucine, leucine, phenylalanine, tryptophan, and tyrosine in the dysbiotic IBD cluster (Figure 2a-b, Table S4b).

Moreover, Ni et al. observed that fecal amino acid concentrations correlated with the abundance of Proteobacteria species and with the severity of the disease (Ni et al., 2017). Based on these findings, Ni et al. proposed that increased proteobacterial utilization of nitrogen for amino acid biosynthesis plays a role in the development of dysbiosis and Crohn’s Disease (Ni et al., 2017). To identify microbes significantly contributing to fecal amino acids, we computed the quantitative microbial contributions to the 22 amino acid metabolites that were present in the reconstructions and measured by Ni et al. in the 108 microbiomes. The proteobacterial contributions to all amino acid metabolites were indeed substantially increased in the dysbiotic cluster (Figure 4b). In contrast, in the healthy controls and the non-dysbiotic IBD cluster, amino acids were mainly synthesized by Bacteroidetes and Firmicutes representatives (Figure 4b).

To elucidate the taxa relevant for amino acid biosynthesis in more detail, the contribution profiles on the strain level were predicted for the examples of glycine, phenylalanine, leucine, tyrosine, and tryptophan (Figure S7-11). In the healthy controls and in the non-dysbiotic cluster, these amino acids were mainly synthesized by commensal bacteria, such as *Alistipes*, *Bacteroides*, *Faecalibacterium*, *Roseburia*, and *Eubacterium rectale* (Figure S9-12). In contrast, mainly opportunistic pathogens, such as *Bacteroides fragilis*, *Escherichia*, *Haemophilus*, *Klebsiella* and *Streptococcus* synthesized amino acids in the dysbiotic cluster (Figure S7-11). In summary, modelling revealed an increased biosynthesis potential for amino acids and increased proteobacterial contributions to amino acids in the microbiomes of patients with dysbiosis, in agreement with the findings of Ni et al. (Ni et al., 2017).

## Discussion

We have systematically profiled the metabolic potential of 108 individual microbiomes *in silico*. The metabolic profiles varied greatly across individual microbiomes reflecting the variation in microbial composition. We determined the net production and uptake potential of each microbiome, the qualitative and quantitative presence of reactions and pathways in each microbiome, the correlations between net production potential and reaction abundance, and finally the quantitative contributions of all strains present in the microbiomes to all secreted metabolites. Our framework resulted in the characterization of the metabolic networks of each microbiome with previously unpreceded detail and scope.

We investigated the computed microbiome properties to identify features that separated dysbiotic IBD microbiomes from non-dysbiotic microbiomes and found that a wide variety of metabolic network properties and fluxes differed between the groups (Table S4b). First, the qualitative and quantitative presence of reactions and pathways differed in dysbiotic microbiomes (Figure 3b, Table S4b). As a result of their disturbed metabolic network structure, dysbiotic microbiomes also demonstrated an altered potential to take up and secrete metabolites (Figure 2, Table S4b). The quantitative contributions of each strain to each secreted metabolite differed clearly between non-dysbiotic and dysbiotic microbiomes (Figure 3c, Figure 4, Table S4b). IBD microbiomes with dysbiosis were characterized by a depletion in contributions of Bacteroidetes and Clostridia and an enrichment of Bacilli and Proteobacteria contributions (Figure 4). This constraint-based modelling framework enabled us link microbes and metabolites in the context of the gut microbial community resulting in detailed biosynthesis profiles of each metabolite (Figure S4-13). We confirmed known microbe-metabolite links, e.g., butyrate production of *Faecalibacterium* and *Roseburia* (Machiels et al., 2014) (Figure S4), and lactate production of *Lactobacillus*, *Streptococcus*, and *Escherichia* (Figure S5). In addition, we propose contributing microbes for less-studied metabolites. For instance, gut microbes regulate circulating tryptophan levels and impaired tryptophan metabolism may play a role in IBD (Agus et al., 2018). We here predict that, e.g., *Alistipes*, *Bacteroides*, and *Odoribacter* can synthesize tryptophan (Figure S11) and thus may play a role in mediating host-microbe interactions in tryptophan metabolism.

Previously, the AGORA resource has been applied to analyze reactions and pathways in the microbiomes of ulcerative colitis and Crohn’s Disease patients’ microbiomes and identify subsystems that were enriched in these groups (Yilmaz et al., 2019). This work already demonstrated that the AGORA reconstructions have useful applications for investigating changes in metabolic pathways as a result of microbial dysbiosis. Here, we similarly took advantage of the curated metabolic networks in AGORA to elucidate how individual microbiomes differ in metabolic network structure. However, in addition to describing changes in microbial community structure in IBD, we also predicted the consequences of the structural changes on the metabolic activity of the microbiomes. This approach resulted in testable hypotheses that could be experimentally validated in subsequent efforts. While one cannot compare fluxes and concentrations (O’Brien et al., 2015), the trends of upregulated and downregulated metabolites in the dysbiotic cluster can nonetheless be compared. Our predictions that the dysbiotic microbiomes had an increased potential to synthesize amino acids mainly due to an increased abundance of Gammaproteobacteria in these microbiomes agreed well with the experimental findings by Ni et al (Ni et al., 2017). Importantly, the microbiome models used in our modelling framework only reflected the influence of altered gut microbial composition in the dysbiotic IBD patients and do not account for host metabolism. This limitation provides the opportunity to distinguish the influence of gut microbial and host metabolism on the host metabolome. Thus, since the models only accounted for microbial metabolic networks and were not confounded by human metabolism or by differences in diet, the modelling also provides evidence that the increased fecal amino acid concentrations measured by Ni et al. are at least in part due to the gut microbiome rather than only of dietary origin.

In future studies, human metabolism could be incorporated into the modelling framework. Since the metabolites and reactions in the models are tractable due to unique, organism-specific identifiers, this will allow us to explicitly predict the contribution of gut microbes and human to each metabolite. The fluxes through microbial and host enzymes of interest could also be traced. Moreover, the personalized models could be contextualized further with individual subjects’ dietary information (Noronha et al., 2019) as well as physiological parameters (Thiele et al., 2018). We have already demonstrated that microbiome models as well as physiological data can be integrated with a whole-body metabolic model of human, enabling personalized, organ-resolved predictions of host metabolic states (Thiele et al., 2018). Such an integrated modelling approach could be used to gain insight into the mechanisms linking altered host metabolism, gut microbiome composition, and gut microbial metabolism in IBD. The whole-body human model also allows the personalized prediction of the urinary, blood and serum metabolomes (Thiele et al., 2018), which have been reported to be altered in IBD patients (Kolho et al., 2017; Schicho et al., 2012; Stephens et al., 2013; Williams et al., 2012). It is known that the blood metabolome contains metabolites of microbial origin and is altered in germfree animals (Wikoff et al., 2009), however, these mammalian-microbial co-metabolites are challenging to trace back to microbial species of origin. Integrated host-microbiome metabolic modelling will allow the personalized prediction of these host metabolomes as a function of dietary input and microbial activity. This will enable the prediction of microbial species contributing to the human blood, urine, and tissue metabolome. Additionally, metabolomic measurements can serve as constraints for the further contextualization of personalized models (Aurich et al., 2016) allowing further insight into the mechanisms behind host-microbe co-metabolism.

In summary, we present an integrative constraint-based modelling framework that enables the comprehensive characterization of personalized gut microbiome models. The framework is scalable, enables the interrogation of gut microbial metabolism on a microbiome-wide scale in cohorts of typical size (>100 individuals), and can be combined with reconstructions of host metabolism. The approach is organism-and molecule-resolved, allows linking metabolites with their reactions of origin, and implicitly accounts for microbe-microbe competition by including organism abundances and a growth-limiting dietary input. The AGORA resource of gut microbe reconstructions (Magnusdottir et al., 2017; Noronha et al., 2019) as well as toolboxes and tutorials for the creation and interrogation of personalized microbiome models (Baldini et al., 2018; Heirendt et al., 2019; Heirendt et al., 2017) are freely available to the scientific community. We envision the computational pipeline described in this study as a powerful resource to facilitate the interpretation of *in vitro* and *in vivo* experiments and to guide the design of prospective studies. Ultimately, an iterative pipeline of computational predictions and experimental validation may yield in the discovery of novel therapeutic and dietary interventions targeting the host-gut microbiome-diet axis.

## Acknowledgements

We thank Dr. Laurent Heirendt and Federico Baldini for assisting in setting up the initial simulations and models, and Dr. Johannes Hertel for statistical advice. We also thank Prof. Dr. Frederic D. Bushman, Prof. Dr. Gary D. Wu, Assistant Prof. Dr. Kyle Bittinger, Prof. Dr. Hongzhe Lee, and Prof. Dr. James D. Lewis for valuable discussions.

## Author Contributions

Conceptualization, A.H. and I.T..; Methodology, A.H.; Formal Analysis, A.H.; Investigation, A.H.; Writing ‒Original Draft, A.H..; Writing ‒Review & Editing, A.H. and I.T.; Visualization, A.H.; Funding Acquisition, I.T.; Supervision, I.T.

## METHODS

### Creation of personalized models

Paired end Illumina raw reads of 83 dysbiotic Crohn’s Disease patients in the PLEASE cohort (Lewis et al., 2015) and of 25 healthy controls in the COMBO cohort (Wu et al., 2011) had been previously retrieved from NCBI SRA under SRA: SRP057027 (Bauer and Thiele, 2018). The reads had been preprocessed and mapped onto the reference set of AGORA genomes (Bauer and Thiele, 2018). Personalized models for the 108 samples were created using Version 1.03 (published on 25.02.2019, available at https://www.vmh.life) of the AGORA resource (Magnusdottir et al., 2017). To build the personalized models, the COBRA Toolbox (Heirendt et al., 2019) extension Microbiome Modelling Toolbox (Baldini et al., 2018) was used. Personalized microbiome models were created using the mgPipe module, as described previously (Heinken et al., 2019). For all strains identified in at least one metagenomics sample, the corresponding AGORA reconstructions, if available, were joined into one global constraint-based microbiome community reconstruction (Baldini et al., 2018). For each of the 108 metagenomic samples, the strain-level abundances that had been previously mapped onto AGORA served as input data for deriving a personalized microbiota model from the global reconstruction, which consisted of the joined AGORA reconstructions corresponding to each strain present in the sample. The flux through each AGORA strain’s model was coupled to its respective biomass objective function. Each personalized model contained a community biomass reaction which was parameterizing by applying the strain-level abundances as stoichiometric values for each microbe biomass reaction in the community biomass reaction. These constraints enforced that all strains grew at the experimentally measured ratios. The models were further contextualized as follows: To simulate a realistic intake of dietary nutrients in mmol per g dry weight per hour in the 108 microbiome models, an Average European diet was retrieved from the Diet Designer resource on the Virtual Metabolic Human website (https://www.vmh.life) (Noronha et al., 2019). The diet was converted to uptake fluxes through a dedicated Microbiome Modelling Toolbox function (*convertVMHDiet2AGORA.m*). Moreover, to account for host metabolism, the uptake of metabolites of host origin known to be present in the intestine (e.g., primary bile acids, host glycans) were allowed using standard COBRA Toolbox functions (Heirendt et al., 2019). Finally, to simulate a realistic turnover of microbial biomass, the allowed flux through the community biomass reaction was set to be between 0.4 and 1 mmol × person^−1^ × day^−1^, corresponding to a fecal emptying of once every three days to once a day.

### Prediction of metabolic profiles

Absolute reaction presence and reaction abundances on the total community, phylum, and genus level were calculated in MATLAB using dedicated Microbiome Modeling Toolbox functions (*calculateReactionPresence.m*, *calculateReactionAbundance.m*). The computation of the total community metabolite production potential, total community metabolite uptake potential, and the contribution of each strain to each metabolite was performed in Julia (https://julialang.org) using the Julia implementation of flux balance analysis, COBRA.jl (Heirendt et al., 2017). COBRA.jl was performed on a high-performance cluster using the IBM CPLEX solver (IBM, Inc.) through the CPLEX interface for Julia. A customized Julia script was used that retrieved all dietary exchange reactions, fecal secretion reactions, and strain-specific internal exchange reactions for each microbiome model. This resulted on average in 13,677 exchange reactions per microbiome model, which were then each minimized and maximized using distributed flux balance analysis (Heirendt et al., 2017). The fluxes were exported from Julia using the customized Julia script and further analyzed in MATLAB.

### QUANTIFICATION AND STATISTICAL ANALYSIS

#### Statistical analysis

Wilcoxon rank sum test, and correction for false positive discovery rate were performed in MATLAB using the ranksum and mafdr functions, respectively. Correlations between computed fluxes and species abundances were calculated using a dedicated Microbiome Modeling Toolbox function (*correlateFluxWithTaxonAbundance.m*).

#### Visualization

Random forests analysis was performed using the online implementation of MetaboAnalyst 4.0 (Chong et al., 2018) (https://www.metaboanalyst.ca). Total metabolite fluxes were plotted using the violinplot implementation in MATLAB (https://github.com/bastibe/Violinplot-Matlab). Phylum to amino acid and subsystem contributions were visualized using the online implementation of Circos (Krzywinski et al., 2009) (http://circos.ca). All other data was visualized in R version 3.5.3 (Team, 2016) (https://www.r-project.org) using the pheatmap and vegan packages.

## DATA AND SOFTWARE AVAILABILITY

The AGORA resource (Magnusdottir et al., 2017) and the Average European diet simulation constraints are freely available at the Virtual Metabolic Human (Noronha et al., 2019) website (https://vmh.life). The COBRA Toolbox (Heirendt et al., 2019), the Microbiome Modeling Toolbox (Baldini et al., 2018), and COBRA.jl (Heirendt et al., 2017) are freely available at GitHub (https://github.com/opencobra). Tutorials to create the sample-specific constrained models and to repeat the analysis performed in this paper are available at https://github.com/opencobra/COBRA.tutorials.

